# Protein Surface Printer for Exploring Protein Domains

**DOI:** 10.1101/2020.07.26.222265

**Authors:** Yang Li, Baofu Qiao, Monica Olvera de la Cruz

## Abstract

The surface of proteins is vital in determining protein functions. Herein, a program, Protein Surface Printer(PSP), is built that performs multiple functions in quantifying protein surface domains. Two proteins, PETase and cytochrome P450, are used to validate that the program supports atomistic simulations with different combinations of programs and force fields. A case study is conducted on the structural analysis of the spike proteins of SARS-CoV-2 and SARS-CoV, and the human cell receptor ACE2. Although the surface domains of both spike proteins are highly similar, their receptor binding domains(RBDs) and the O-linked glycan domains are structurally different. Statistically, the outer surface of ACE2 displays less correlation with the RBD of SARS-CoV-2 than that of SARS-CoV. The O-linked glycan domain of SARS-CoV-2 is highly positively charged, which may promote binding to negatively charged human cells. Our program paves the way for an accurate understanding of protein binding for aggregation and ligand recognition.

## 1. Introduction

Surface domains of a protein are defined as the areas where amino acids with similar polarity are distributed close to each other, which collectively impact protein interactions including protein-protein interactions and ligand binding by forming a domain-like structure on protein surfaces. Though many previous studies have unveiled the importance of protein surfaces in functions such as protein adsorption^1^ and molecule recognition, ^2^ little research has been done to demonstrate the role of surface domains in those processes. Recently, protein surface domains were found to be able to selectively bind amphiphilic random copolymers in foreign environments.^3,4^ Specifically, the polar protein surface domains were found to selectively bind the polar components of the random copolymers, with the nonpolar components of the random copolymers extending into the organic environment. The formed protein-polymer complexes thus display a core (protein) - shell (polymer) reverse micellar structure, which consequently preserved the enzymatic activity of proteins. Furthermore, protein surface domains can distinctly orient water neighbors at the protein surface.^5^ For instance, negatively charged surface amino acids can orient water dipoles towards the proteins, whereas positively charged surface amino acids can orient water dipoles against the proteins. Moreover, different types of surface amino acids demonstrate different capabilities in orienting water neighbors and in influencing water dynamics. Consequently, the water shell near the protein surface forms an alternative pattern according to the protein surface domains.^5^ The protein local hydration behavior is expected to impact protein solubility and assembly,^6–8^ enzymatic substrate hydrolysis,^9,10^ and ligand recognition. ^6,11^

Given the crucial role of protein surface domains and lack of related studies, it is of paramount importance to quantitatively characterize the protein surface domains and consequently their impacts on protein functions. For instance, by integrating machine learning techniques^12^ with protein surface domain information, it may be feasible to predict protein solubility, protein-protein interactions and protein-ligand binding affinity. Currently, there is well-developed software such as VMD^13^ that can visualize proteins and simulation tra-jectories. However, functions to quantity protein surface domains are limited. It is hard to obtain protein surface information by simple observations of the proteins alone. Furthermore, multiple molecular dynamic (MD) software is prevailing but they produce different output formats, most of which are not processable by other software. The obstacle limits the analysis of results from the MD simulation results if researchers are not familiar with the software.

Inspired by the previous works, we have developed a program, known as Protein Surface Printer (PSP), to analyze protein surface domains and print surface domain information. The program was developed with a Python package MDAnalysis^14,15^ to perform multiple functions 1) analyze MD simulation trajectories in the formats of packages GROMACS,^16^ NAMD,^17^ AMBER^18^ and OpenMM^19^ using different atomistic force fields of AMBER,^20^ CHARMM^21^ and AMOEBA,^22,23^ 2) process a single frame or multiple frames, 3) read and clip trajectories, 4) align proteins for multiframe trajectories, 5) collect and print surface domain information, 6) generate files containing protein surface domain information, 7) create files that can be visualized by software (such as VMD).

Two proteins, PETase and cytochrome P450, were used as examples to test the validity of the new program PSP. PETase is an efficient poly(ethylene terephthalate) plastic-degrading enzyme.^9^ Cytochrome P450 is one of the most versatile biocatalysts in nature. ^24,25^ Both proteins are crucial in a broad variety of chemical reactions particularly important for pharmaceuticals, terpenes, gaseous alkanes, etc.^24^ Around 75% of drugs are metabolized by P450 proteins.^26^ After validation, a special case study was conducted on the spike proteins of the recent SARS-CoV-2 virus and its close relative SARS-CoV.

## 2. Methodology

### 2.1 Protein Surface Printer (PSP)

Surface residues are defined as residues that are in contact with solvent atoms. Some protein molecules are not solid inside and small solvent molecules (such as water molecules) can diffuse inside the protein. Thus, firstly, PSP excludes solvent atoms that are inside the proteins by finding the largest solvent cluster around the protein. Secondly, PSP calculates the distance between an amino acid and its nearest solvent atom. If the distance is less than 3.5 Å, which is the hydrogen donor-acceptor length, ^27^ the amino acid will be labeled as a surface residue. If the simulation trajectory is provided as the input to the program PSP, it will analyze the whole trajectory, which is termed as multi-frame analysis here. Otherwise, the program will only analyze one individual configuration, termed as single-frame analysis here. Lastly, PSP will print out surface domain information depending on user-defined options. The running logic is shown in Scheme 1.

PSP categorizes surface residues into 4 kinds based on their charges and polarities as shown in Table 1.^5,13^

**Table 1:** Surface domain categories.

| Surface domain | Residues |
| --- | --- |
| Positively charged | ARG; HSP (charged HIS); LYS |
| Negatively charged | ASP; GLU |
| Polar neutral | ASN; CYS; GLN; SER; THR; TYR; GLY; neutral HIS |
| Nonpolar | ALA; LEU; VAL; ILE; PRO; PHE; MET; TRP; |

### 2.2 PETase and P450

PETase was used to test the multi-frame function. In a previous work,^5^ we equilibrated PETase (+6e), Cl^-^ counterions, and 0.1 M salt of Na^+^ and Cl^-^ in a water box for 300 ns. Here we extended the simulations by another 2 ns using a variety of MD simulation programs (GROMACS,^16^ NAMD,^17^ AMBER^18^ and OpenMM^19^) and atomistic simulation force fields (CHARMM36m, ^21^ CHARMM36,^28^ AMBER14SB^20^ and AMOEBA 13^23^). The isothermal-isobaric ensemble (constant number of particles, temperature and pressure, NTP) was applied to all the simulations. The other parameters are listed in Table 2 and the details can be found in the references therein. The structure of cytochrome P450 was downloaded from the Protein Database with the PDB ID 1BU7^29^ and was dissolved in water with Gromacs. The resulting file was used for a single frame analysis.

**Scheme 1:**
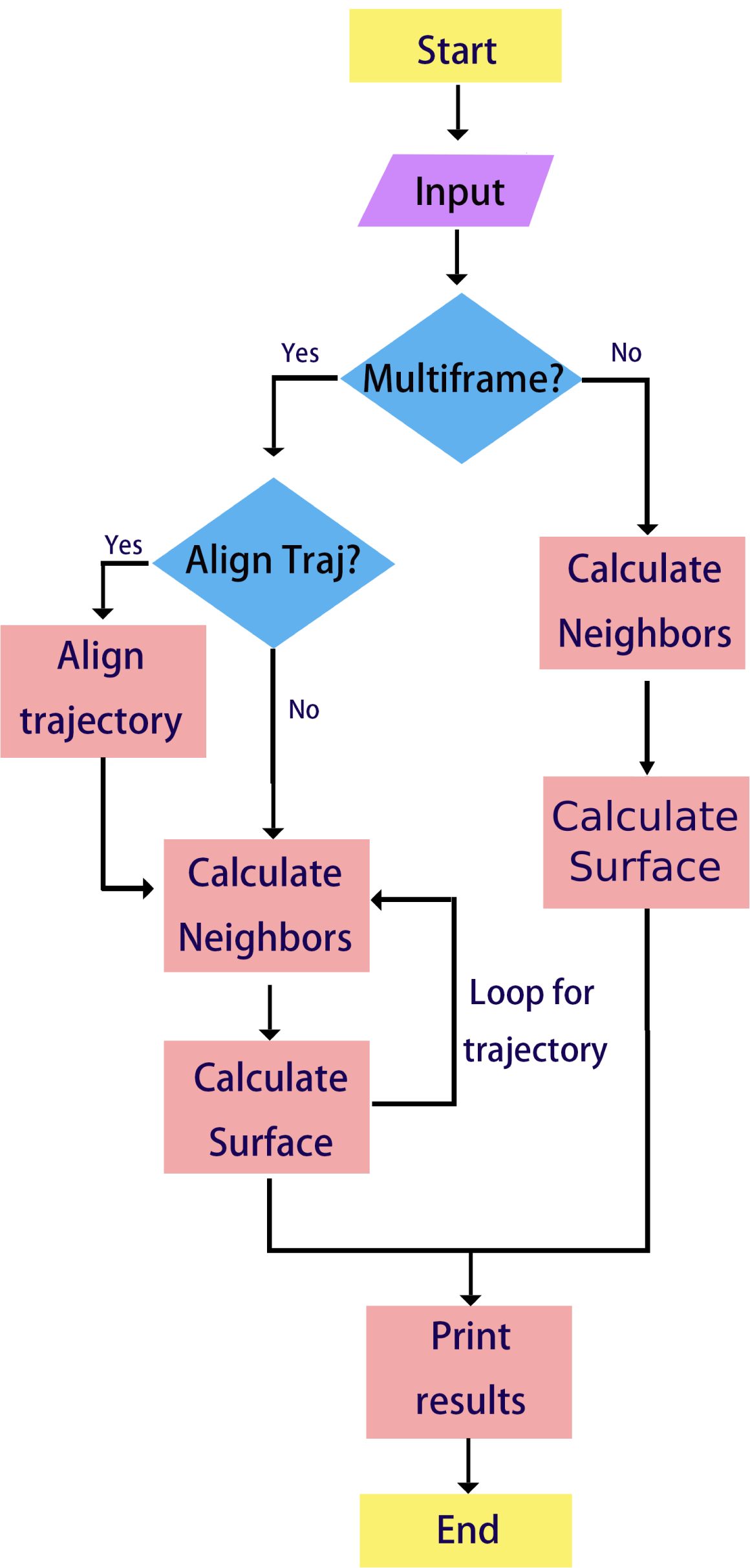
Flow chart of Protein Surface Printer running progress.

**Table 2:**
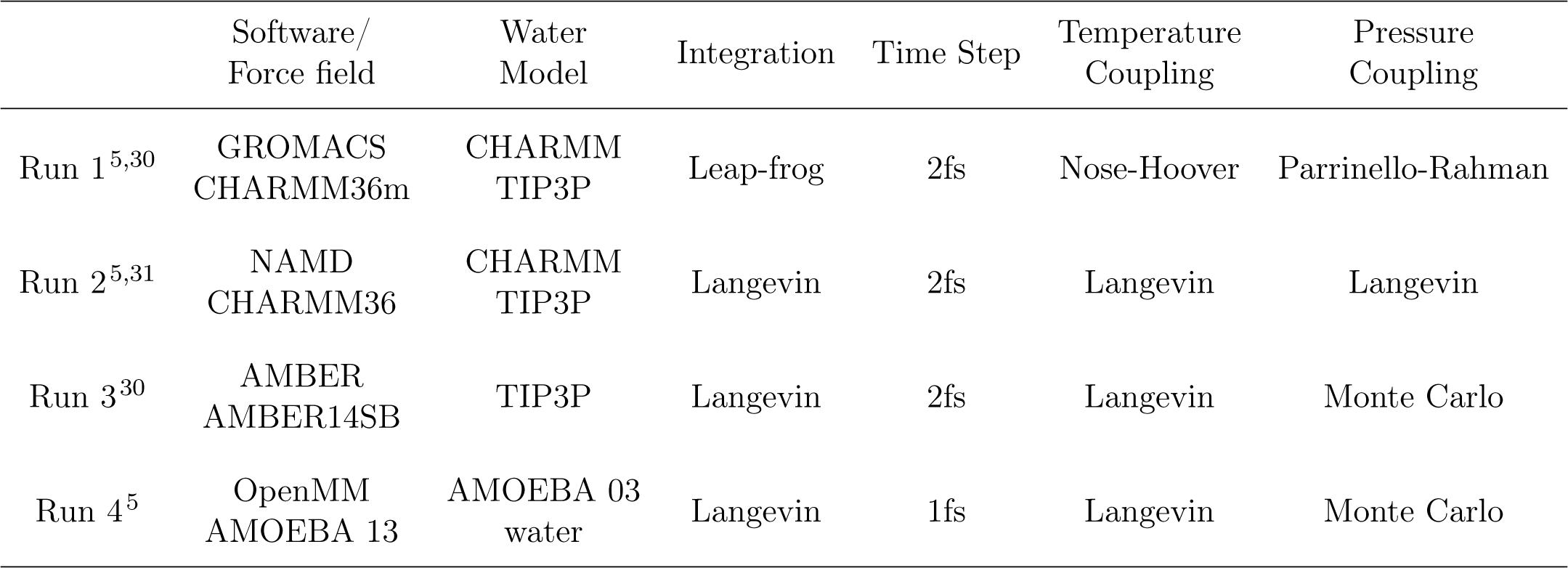
Parameters of the MD simulations on PETase.

|  | Software/<br>Force field | Water<br>Model | Integration | Time Step | Temperature<br>Coupling | Pressure<br>Coupling |
| --- | --- | --- | --- | --- | --- | --- |
| Run 1<sup>5,30</sup> | GROMACS<br>CHARMM36m | CHARMM<br>TIP3P | Leap-frog | 2fs | Nose-Hoover | Parrinello-Rahman |
| Run 2<sup>5,31</sup> | NAMD<br>CHARMM36 | CHARMM<br>TIP3P | Langevin | 2fs | Langevin | Langevin |
| Run 3<sup>30</sup> | AMBER<br>AMBER14SB | TIP3P | Langevin | 2fs | Langevin | Monte Carlo |
| Run 4<sup>5</sup> | OpenMM<br>AMOEBA 13 | AMOEBA 03<br>water | Langevin | 1fs | Langevin | Monte Carlo |

### 2.3 SARS-CoV-2 and SARS-CoV

Here the NCBI GenBank access code MN908947 (protein identification number QHD43416.1) is employed for SARS-CoV-2^32^ and the NCBI GenBank access code AY278741 (protein identification number AAP13441.1) is employed for SARS-CoV. ^33^ The sequences are provided in the Supporting Information. The sequences were first uploaded to the SwissModel server.^34^ The experimental structure (PDB ID: 6VSB^35^) was found to display 99.26% identity to the spike protein of SARS-CoV-2, and the experimental structure (PDB ID 6ACD^36^) demonstrates 100% identity to the spike protein of SARS-CoV. These two protein structures were then employed to build the homology structure model of SARS-CoV-2 and SARS-CoV. The models are provided in the Supporting Information as 6VSB_model.pdb and 6ACD_model.pdb, respectively.

## 3. Results

### 3.1 Multiple Frames Analysis with Trajectory Alignment

PETase was used as an example to test the multi-frame function and the trajectory alignment function. PETase is positively charged with an overall charge of +6e with e being the elementary charge. PETase is roughly of spherical shape, which is ideal for testing the program.

We carried out MD simulations using different combinations of MD simulation packages and force fields: Gromacs/CHARMM36m, AMBER/AMBER14SB, NAMD/CHARMM36 and OpenMM/AMOEBA 13. The protein surface domain distributions are provided in Fig. 1(a-d), respectively. Besides the legends shown in Fig. 1, the extent of opacity indicates the distance between the residues and the center of mass or center of geometry of the protein depending on the input setting. The more transparent a dot is, the closer the residue represented by the dot locates to the center of the protein. The obtained distributions are highly similar to each other for all four types of protein surface domains. When superimposed, the patterns of surface domains are almost the same. We also calculated the average probability and standard deviation for each type of protein surface domain, which are presented in Fig. 1e. Again a high similarity is observed. These calculations thus demonstrate that the PSP program supports a variety of MD simulation packages and force fields, which are extensively employed for protein simulations. The program is expected to support other types of MD simulation trajectories and force fields that are compatible with MDanalysis.

**Fig. 1:**
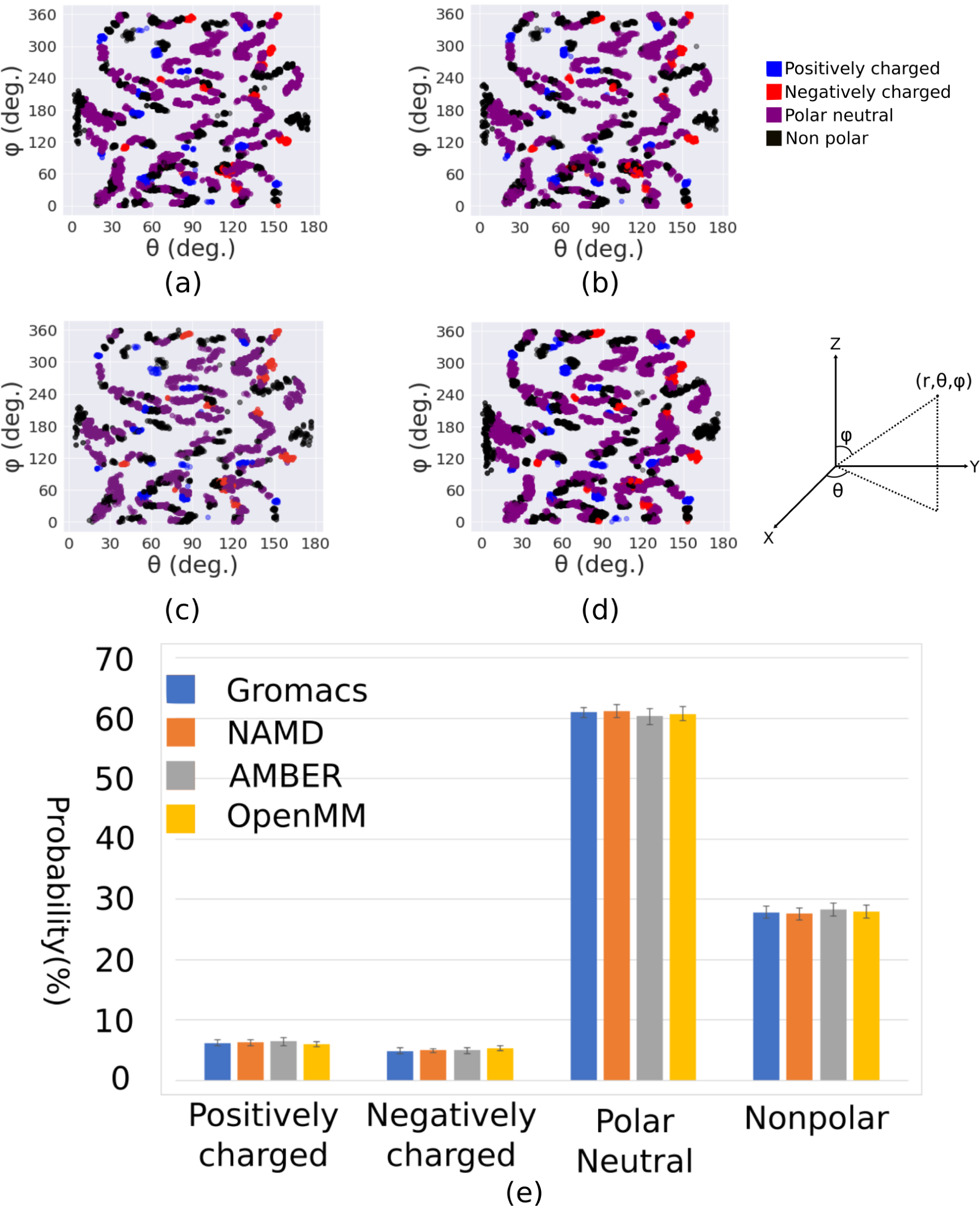
Protein surface domains obtained from (a) Gromacs/CHARMM36m (b) NAMD/CHARMM36 (c) AMBER/AMBER14SB (d) OpenMM/AMOEBA 13. The spherical coordinates θ and φ were employed. (e) The average probability for all types of protein surface domains. Detailed numbers can be found in Supporting Information table S.1.

### 3.2 Single Frame Analysis

P450 is a negatively-charged protein. Each monomer of the P450 dimer (PDB ID: 1BU7) carries a net charge of -15e (including one heme cofactor).^37^ In Fig. 2, the protein surface pattern has two parts, which are roughly symmetric. This is ascribed to the mirror-like orientation of the two subunits of the P450 dimer. Note that in our previous work,^5,37^ only the surface of one of the two subunits was analyzed. In comparison with the corresponding results for a P450 monomer, the probability of nonpolar surface amino acids increases for the P450 dimer, whereas it drops for the other types of surface amino acids. This suggests that the contact region of the two P450 subunits is predominantly composed of polar amino acids, which has been previously observed.^37^

**Fig. 2:**
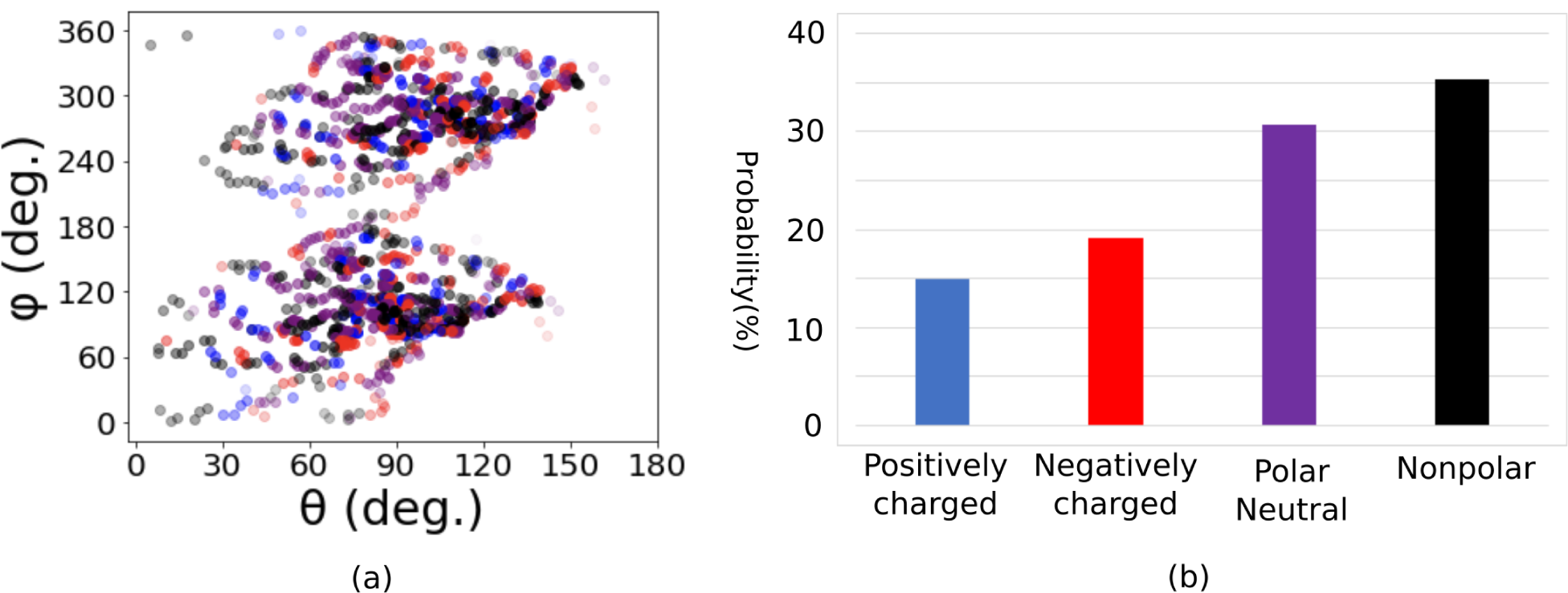
Singe frame surface analysis of P450 with pdb ID 1UB7. Legends are same as those of Fig. 1. (a) Spacial distributions of domains (b) percentages of all kinds of surface domains. Detailed numbers can be found in Supporting Information table S.2.

### 3.3 Quantifying the surface of the spike proteins of SARS-CoV-2 and SARS-CoV, and ACE2

The SARS-CoV-2 has spread to hundreds of countries since its outbreak and infected more than 2,920,000 people at the time of this study. The WHO rated SARS-CoV-2 as *Pandemic*. Early gene study indicated that SARS-CoV-2 (also referred to as 2019-nCoV) is very similar to SARS-CoV.^38^ They both use a densely glycosylated spike protein to gain entry into host cells. The spike protein is a trimeric class I fusion protein that exists in a metastable prefusion conformation that undergoes a substantial structural rearrangement to fuse the viral membrane with the host cell membrane.^35^ It is generally believed that the binding between the spike proteins and the human receptor angiotensin I converting enzyme 2 (ACE2) plays a crucial role in cell fusion. In the present work, we are analyzing the surface features of the spike proteins of SARS-CoV-2 and SARS-CoV, their receptor binding domains and O-linked glycan residues (Fig. 3),^35^ as well as the human ACE2, which is expected to aid the molecular understanding of the binding behavior of SARS-CoV-2 and SARS-CoV to the human cell membrane.

**Fig. 3:**
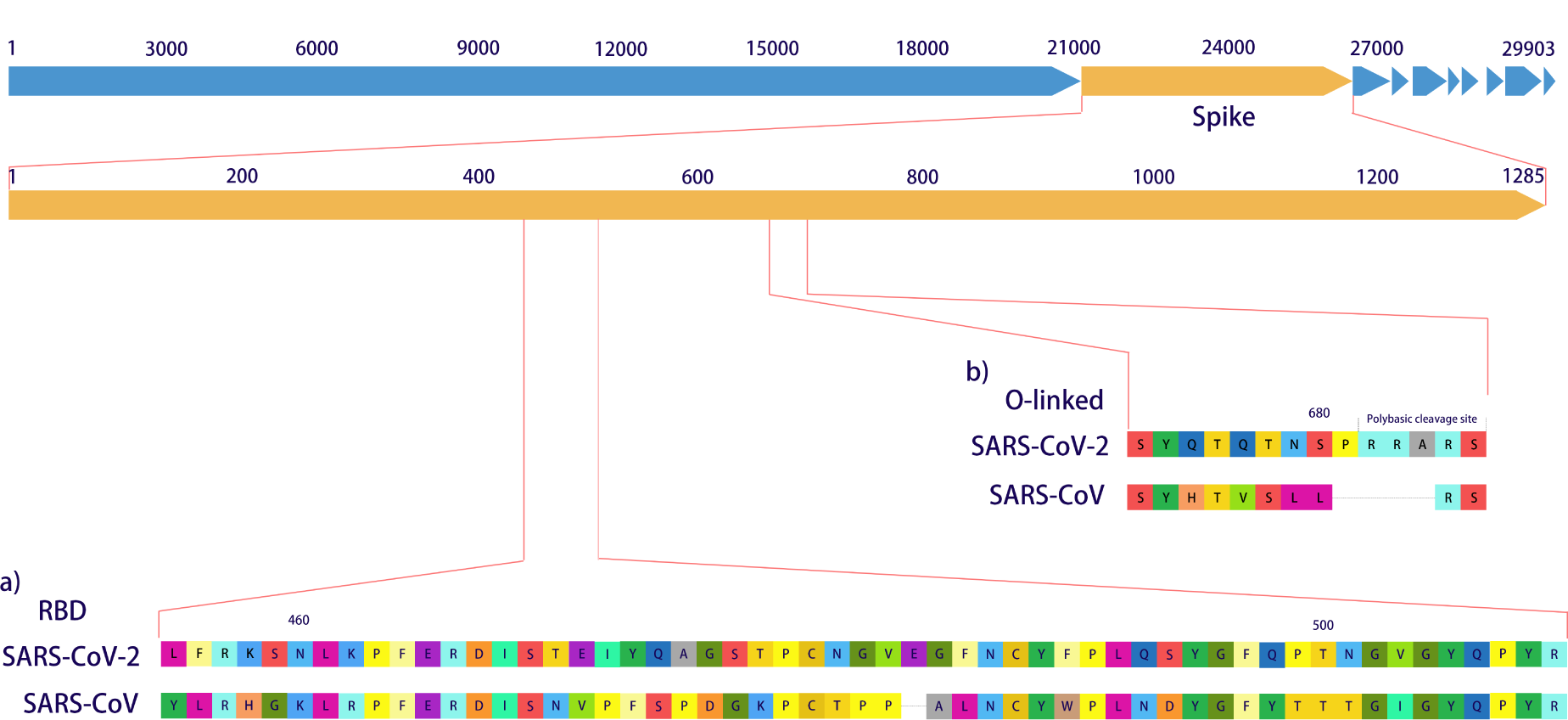
Sequence of (a) receptor binding domains and (b) O-linked residues.

#### 3.3.1 Spike Protein Surface Domains

The homology structures were analyzed using PSP, and the obtained protein surface features are demonstrated in Fig. 4. The compositions of both spike proteins display high similarity. Both proteins are negatively charged: their surface amino acids are 10% - 11% negatively charged and 9% positively charged. The overall negative surface charge has also been experimentally observed for the inner leaflets of plasma membranes^39^ and a broad variety of cancer cells.^40^ A minor difference lies in the polar charge-neutral amino acids at the surfaces. The SARS-CoV-2 has a slightly higher probability (47%) of polar charge-neutral surface amino acids than SARS-CoV (45%). Two spike proteins cannot be distinguished solely by the composition.

**Fig. 4:**
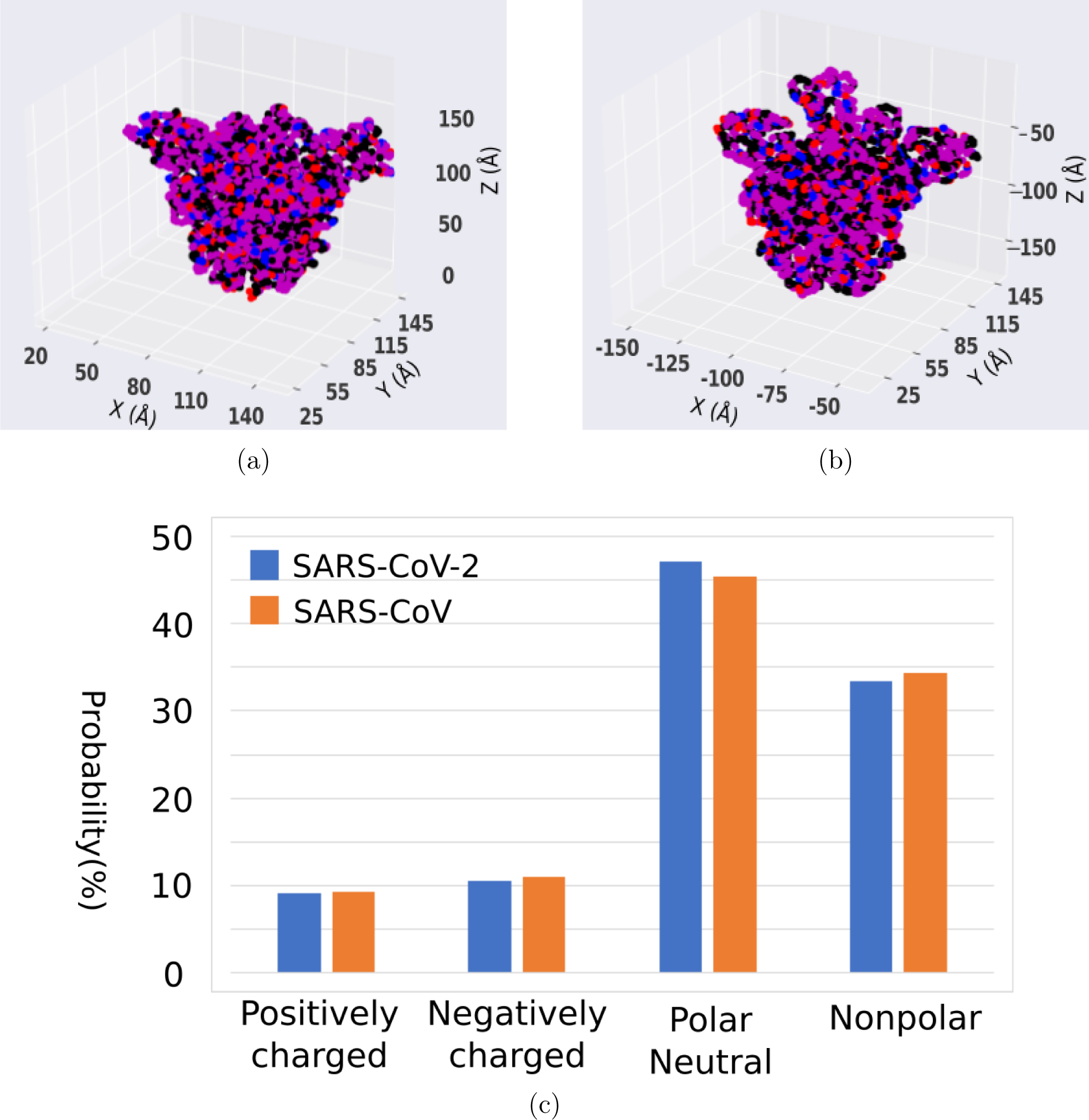
Protein surface domains in Cartesian Coordinates. (a) Model SARS-CoV-2 based on PDB ID 6VSB, (b) model SARS-CoV based on PDB ID 6ACD, and (c) probability of protein surface domains for the spike proteins. Detailed numbers can be found in Supporting Information table S.3.

#### 3.3.2 Receptor Binding Domain (RBD) and ACE2

Early research^41^ suggests that SARS-CoV-2 probably uses the SARS-CoV receptor ACE2 as a cellular entry receptor. Subsequently, more and more research is targeting the binding behavior of SARS-CoV-2 and ACE2. ^42,43^ The receptor binding domains of both spike proteins (Fig. 5) are analyzed with the PSP program developed here. Surprisingly, a significant difference was observed. The SARS-CoV-2 is remarkably less charged with an overall probability of 18% of charged surface amino acids when compared to 26% for SARS-CoV. The difference is predominantly ascribed to a smaller number of negatively charged amino acids for SARS-CoV-2 (10%) than for SARS-CoV (17%). SARS-CoV-2 has more polar neutral residues on its surface (50%) than SARS-CoV (39%). As shown in Fig. 6, analysis of the outer surface of ACE2 supports that it is oppositely charged: around 20% of ACE2 outer surface is negatively charged, which surpasses the probability of positively charged surface amino acids of 10%. The oppositely charged feature between ACE2 and the receptor binding domains of both spike proteins favor the binding of the virus and human cell membranes, which is required for cell entry.

**Fig. 5:**
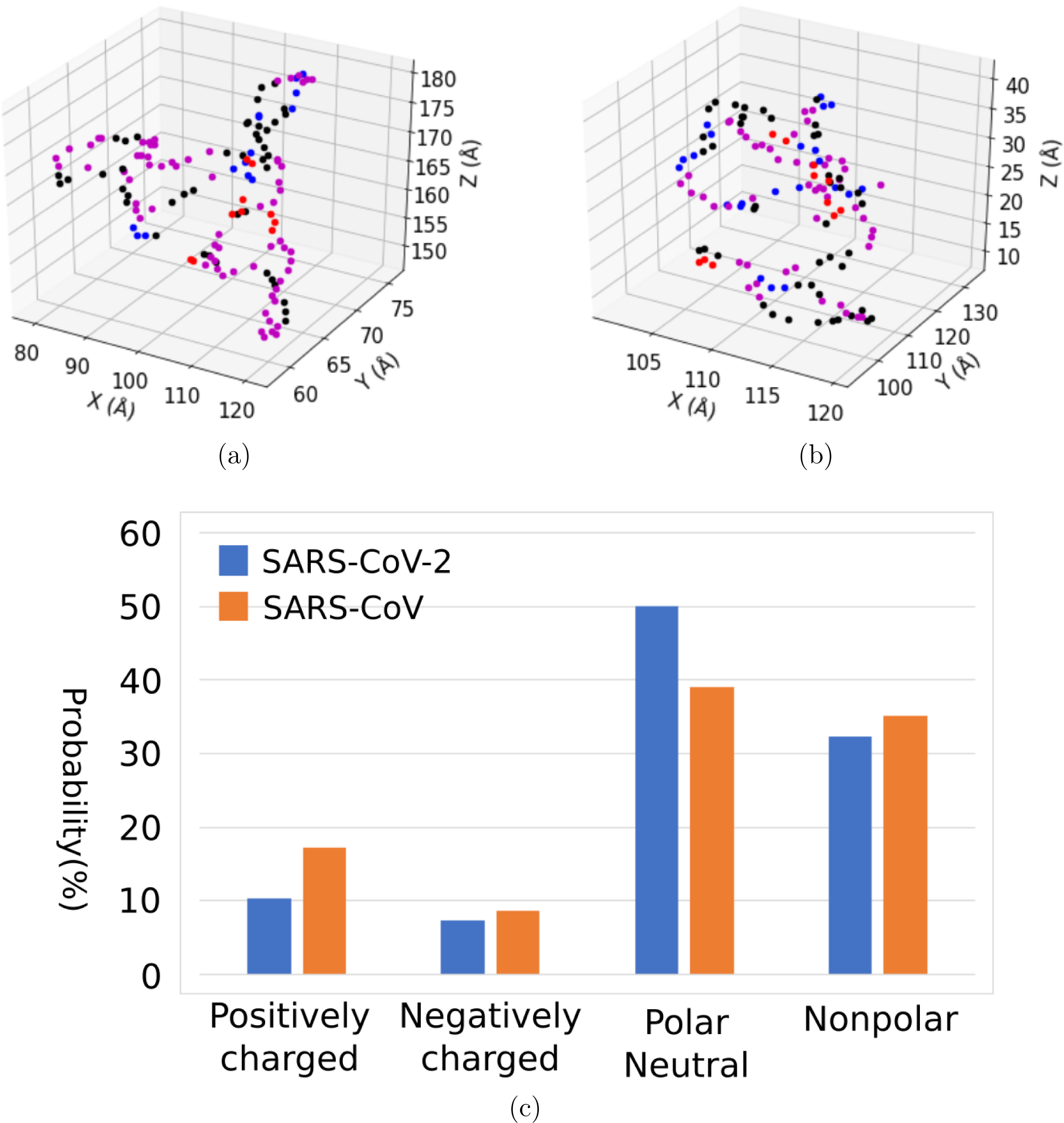
Protein surface domains in Cartesian Coordinates of the receptor binding domains of (a) SARS-CoV-2, (b) SARS-CoV, and (c) probability of each domain. Color legends are the same as those in Figure 1. Detailed numbers can be found in Supporting Information table S.4.

**Fig. 6:**
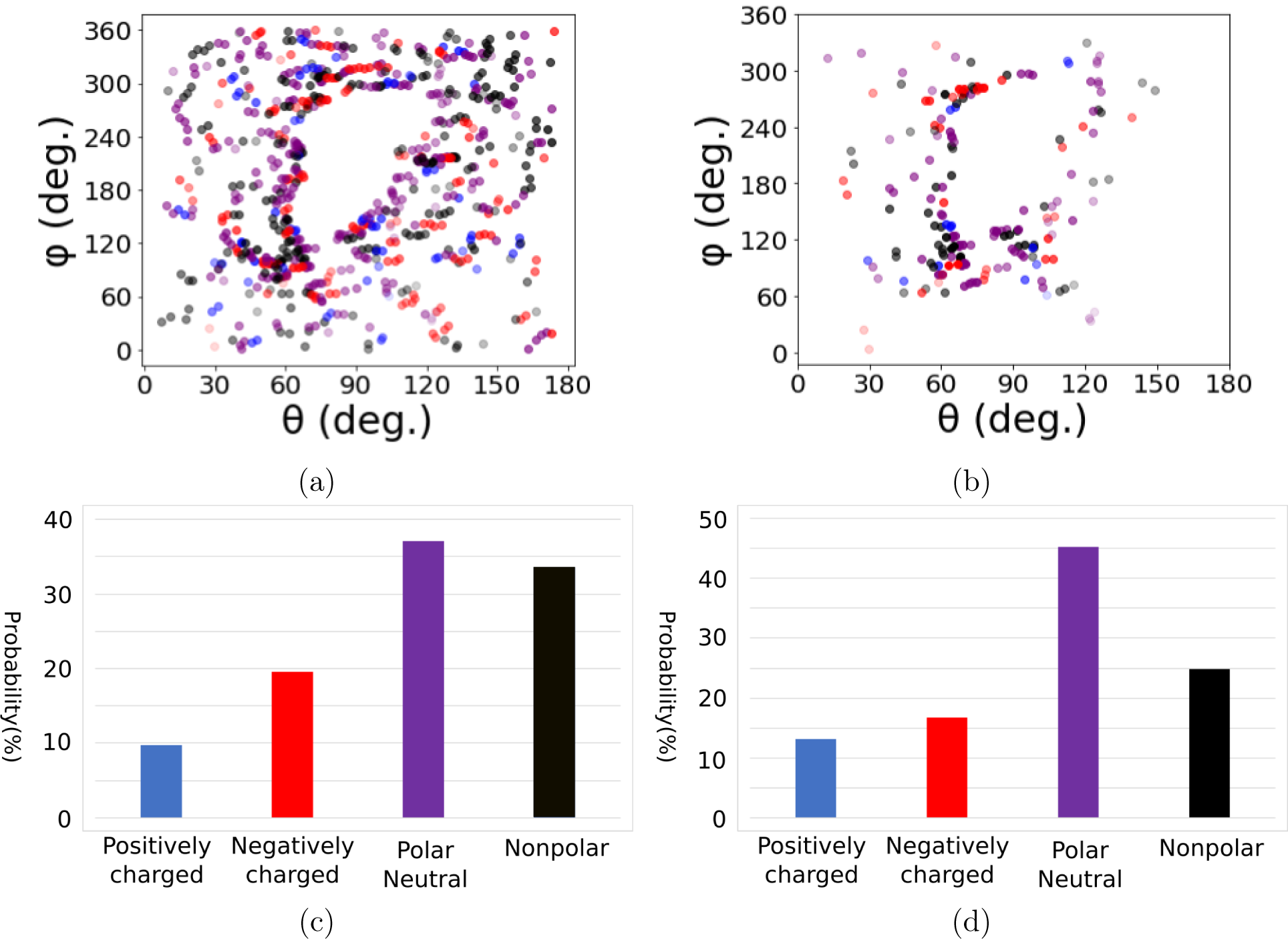
ACE2 surface domains in spherical coordinates. Color legends are the same as those in Figure 1. (a) Outer surface, (b) Inner surface, and Surface domain probability of (c) outer and (d) inner surfaces. Detailed numbers can be found in Supporting Information table S.5.

Note that ACE2 has a concave structure such that it is hard to simply distinguish the surface. Thus, the protein is analyzed separately, with the concave structure as the inner surface and other parts as the outer surface. Remarkably, it is found that the surface domain distributions of the outer surface of ACE2 display strong correlation with the surface domain distributions of SARS-CoV receptor binding domain: 9% positively charged for the former vs 9% negatively charged for the latter; 19% negatively charged for the former vs 17% positively charged for the latter; and similar probabilities of polar charge-neutral and nonpolar surface amino acids for both ACE2 and the receptor binding domain of SARS-CoV. In contrast, the surface domain distributions of SARS-CoV-2 receptor binding domain are less correlated with the surface domain distributions of ACE2 outer surface. The results suggest that based solely on probability, SARS-CoV-2 may not be ideal for binding human ACE2 in contrast to SARS-CoV. These findings are consistent with a prediction based on the structural analysis of the contact region between the two spike proteins and ACE2. ^44^ However, experimental data revealed that RBD of SARS-CoV-2 has a higher affinity than that of SARS-CoV. ^41^ The contradictory result suggests that the high transmissivity of SARS-CoV-2 may not rely solely on the receptor binding domains. For instance, the influence of O-linked glycan residues remains elusive.

#### 3.3.3 O-linked Glycan residues

A unique feature of SARS-CoV-2 is the O-linked glycan residues.^38^ Compared with SARS-CoV, SARS-CoV-2 has four extra residues PRRA located on sequence 681-684, where P, R and A stands for proline, arginine and alanine, respectively. Residues RRAR (sequence 682-685) form a polybasic cleavage site. The functionality of the cleavage site in SARS-CoV-2 remains unclear but is suspected to increase its transmissibility and pathogenesis since the development of the cleavage site was observed with increasing transmissibility of other coronaviruses.^38^ As illustrated in Fig. 7, the inserted residues form a continuous positively charged surface domain. Given the fact that arginine is positively charged and most cell membranes possess negative charges, arginine has previously been shown to improve the cellular uptake of peptides, which is promising for therapeutics applications.^45^ And arginine has been investigated to enhance drug delivery across the blood-brain barrier for Alzheimer’s disease.^46^ It may be hypothesized that in addition to the spike protein-ACE2 binding, the O-liked glycan residues may promote binding of the SARS-CoV-2 with cell membranes, leading to fast cell entry. Mutating arginine to charge neutral or negatively charged amino acids could further illustrate the role of the O-linked glycan residue in immunoevasion.

**Fig. 7:**
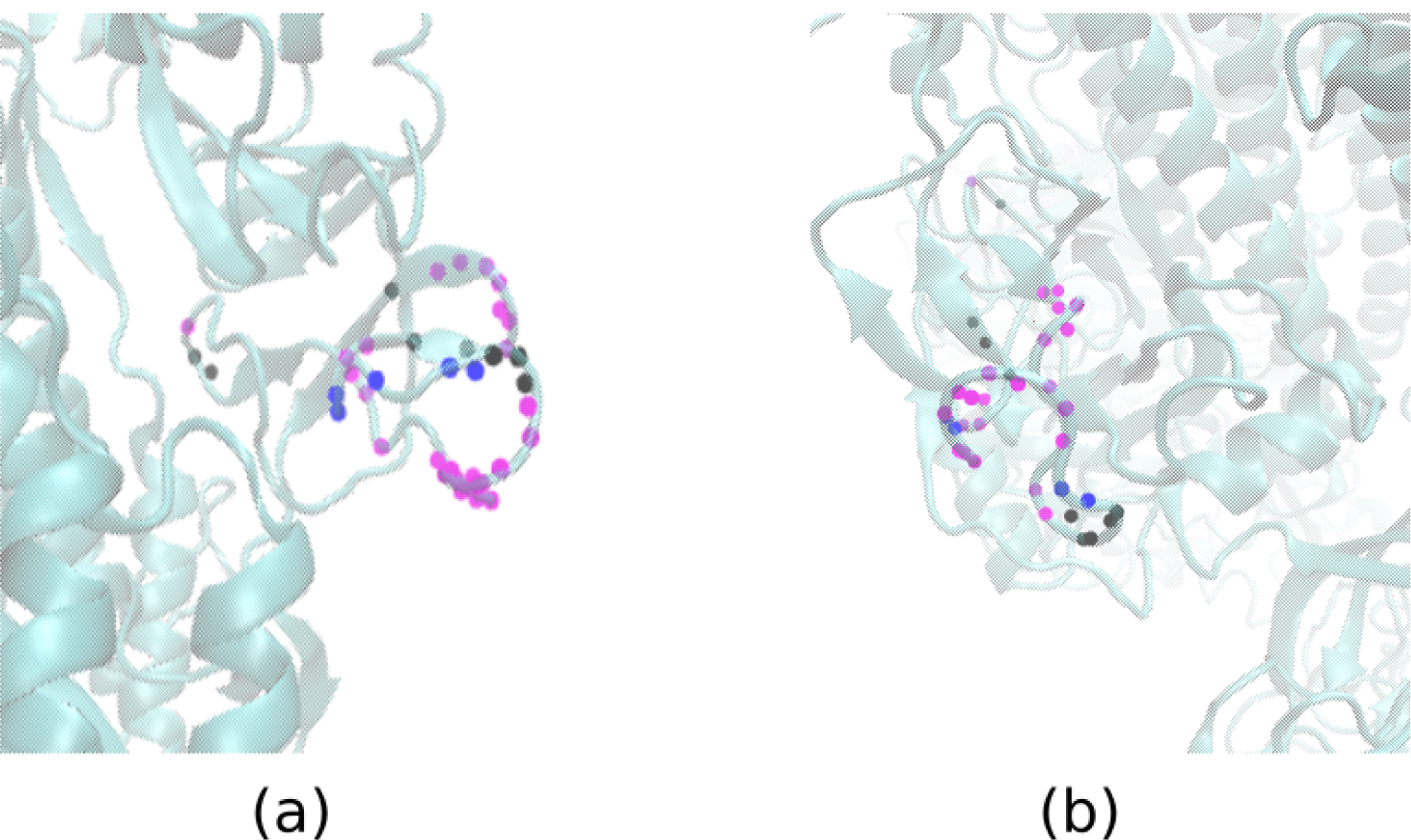
O-like glycan residues. (a) SARS-CoV-2 (b) SARS-CoV. Purple: polar neutral; Blue: positively charged; Red: negatively charged; Black: non-polar.

## 4. Conclusion

A program, PSP, was developed which performs a quantitative investigation of protein surface domains from MD simulation trajectory and single-frame configuration. An analysis of the spike proteins of SARS-CoV-2 and SARS-CoV demonstrates that the two spike proteins are very similar in surface composition in terms of charge and polarity. The largest difference is only 1.5% on polar neutral residues. The receptor binding domain of the new virus SARS-CoV-2 is dominated by polar neutral amino acids (50%), while SARS-CoV consists of comparative amounts of polar neutral residues (39%) and nonpolar residues (35%). An analysis of the outer surface of human cell receptor ACE2 predicts that SARS-CoV-2 may not interact with ACE2 as strongly as SARS-CoV does, which is in agreement with previous structural analysis,^44^ but in contradiction with experimental findings.^41^ This suggests that the high affinity of SARS-CoV-2 may not be exclusively caused by the composition of the receptor binding domains. The highly positively charged feature of the O-linked glycan residues on SARS-CoV-2 may promote binding to the negatively charged human cells. These findings on the SARS-CoV-2 spike protein may provide some clues of value to researchers working on drug discovery and viral pathogenesis.

## Acknowledgments

This work was supported by the U.S. Department of Energy (DOE), Office of Science, Office of Basic Energy Sciences, under Award No. DE-FG02-08ER46539, the Sherman Fairchild Foundation, and the Center for Computation and Theory of Soft Materials at Northwestern University.

## Supporting Information Available

The following files are available free of charge.

Tables with the detailed distribution of surface domains of PETase, P450, the spike proteins of SARS-CoV-2 and SARS-CoV and their receptor binding domains, and the human ACE2. Amino acid sequences of the spike proteins of SARS-CoV-2 and SARS-CoV. 3D print-out of the surface domains. The model protein (6VSB_model.pdb) for SARS-CoV-2 and the model protein (6ACD_model.pdb) for SARS-CoV. The program ProteinSurfacePrinter.py

## Graphical TOC Entry

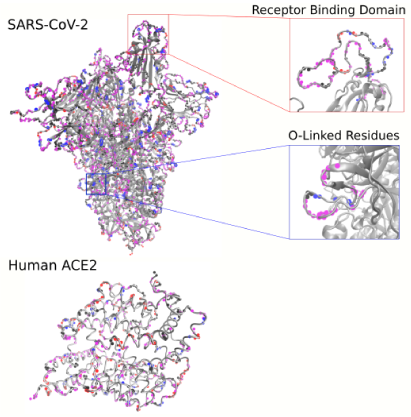

The paper first focuses on the building logic of the program Protein Surface Printer and the validation of the program. Subsequently, the program was utilized to analyze the spikeproteins of the SARS-CoV-2 and its close relative SARS-CoV along with their human binding receptor ACE2.

