## Supplementary figures and images for "Protein Surface Printer for Exploring Protein Domains"

### Supporting Information Movie S.1

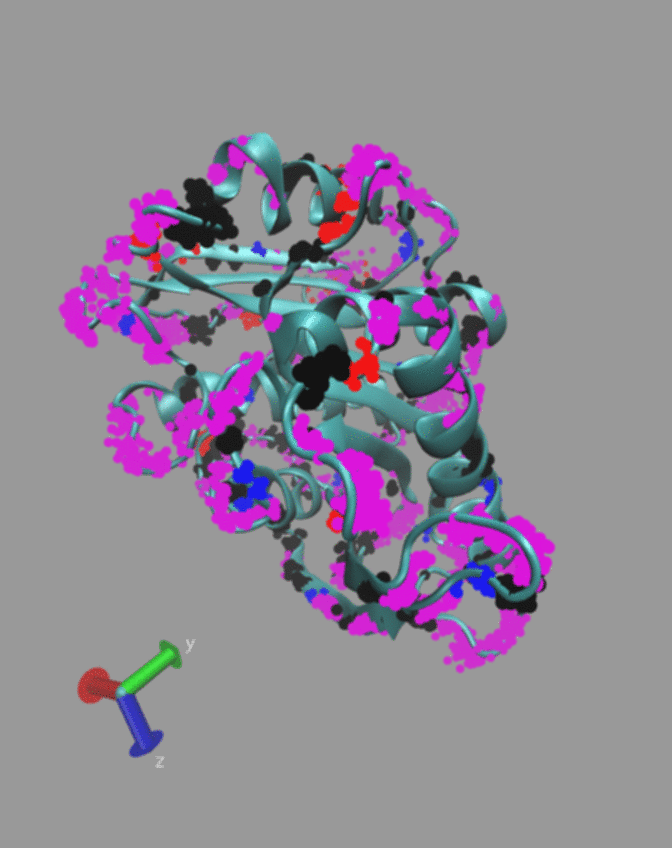

### Supporting Information Movie S.2

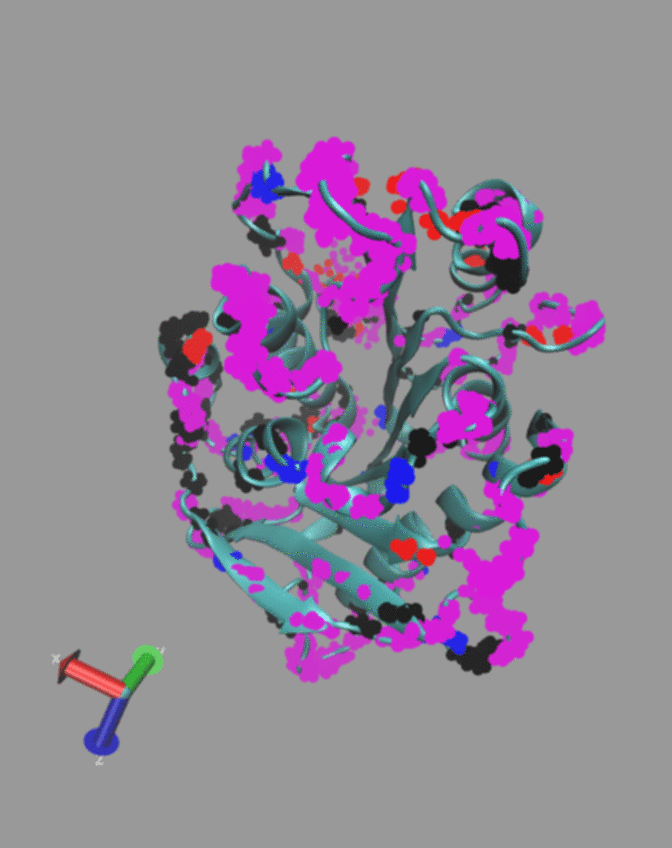

### Supporting Information Movie S.3

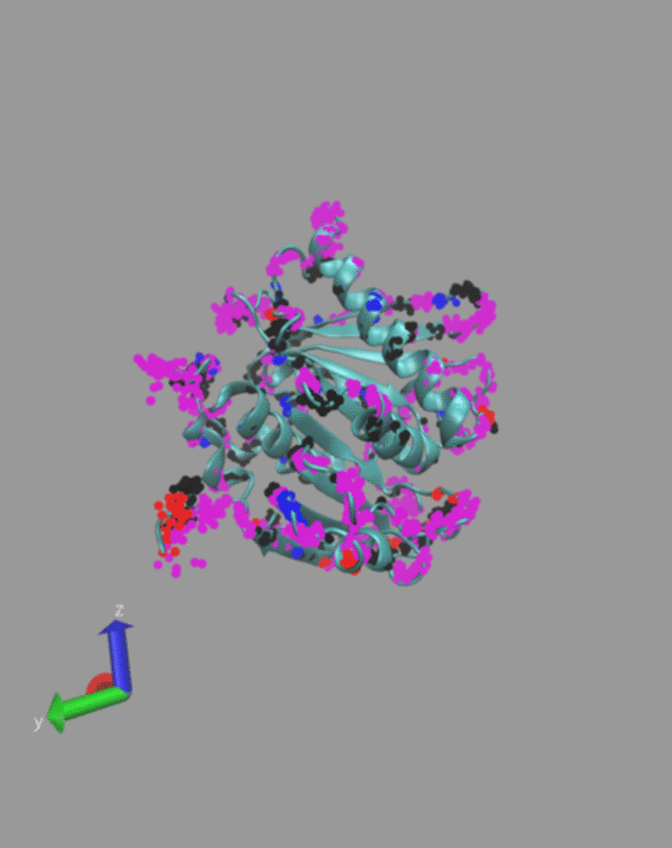

### Supporting Information Movie S.4

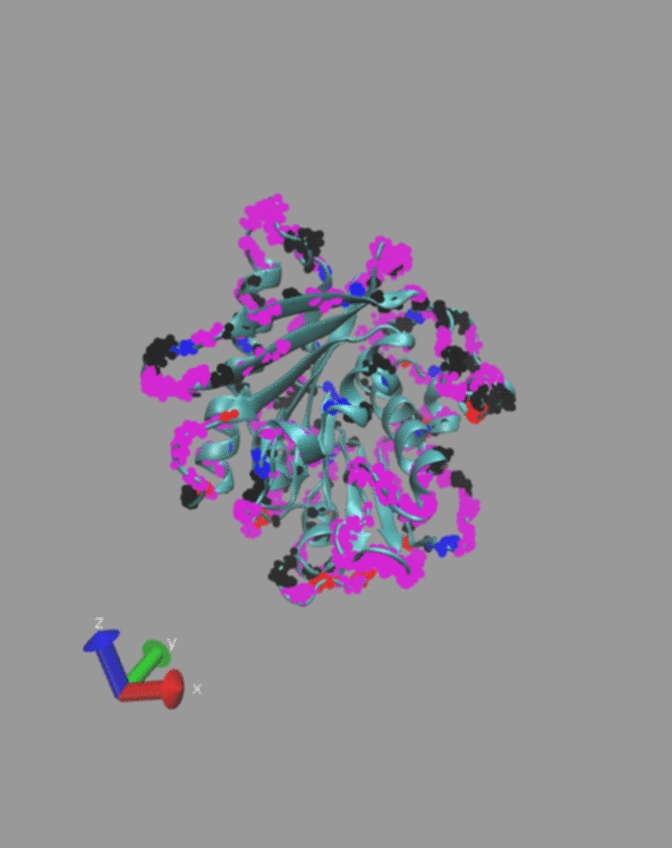

### Supporting Information Movie S.5

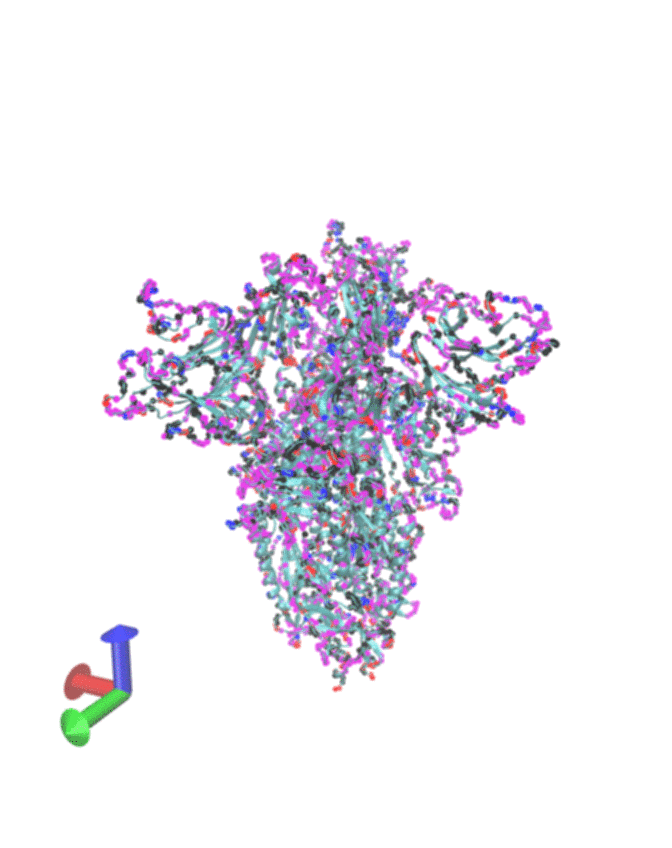

### Supporting Information Movie S.6

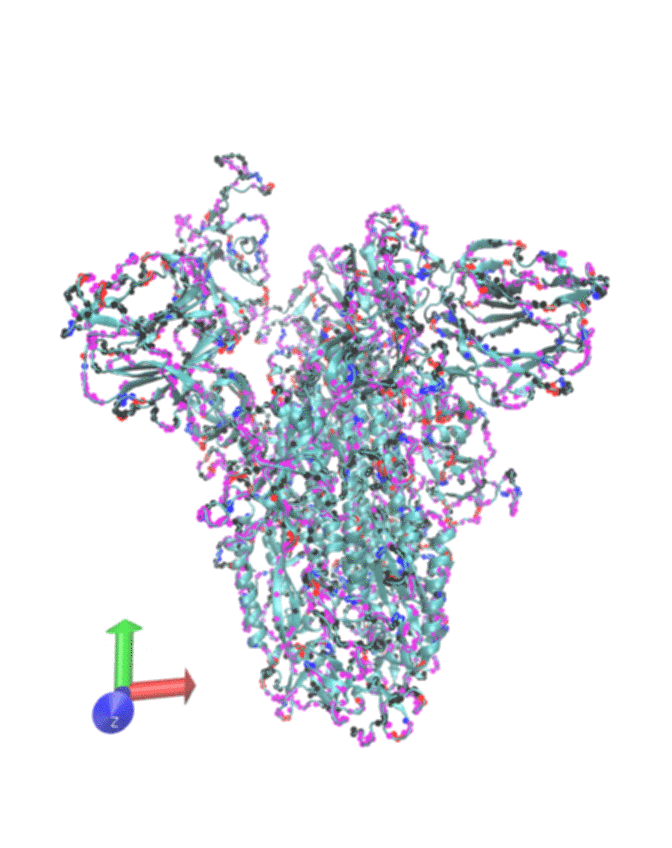

### Supporting Information Movie S.7

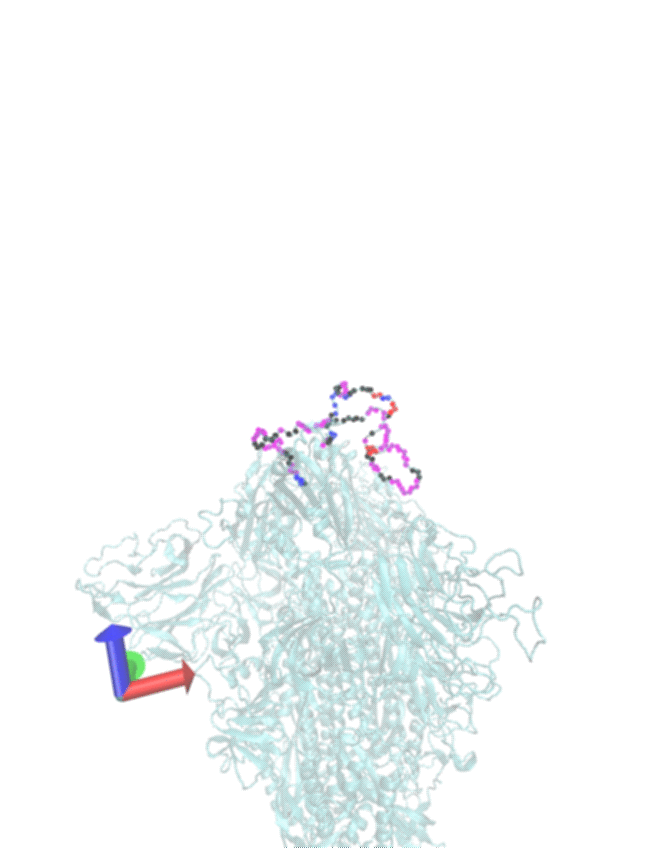

### Supporting Information Movie S.8

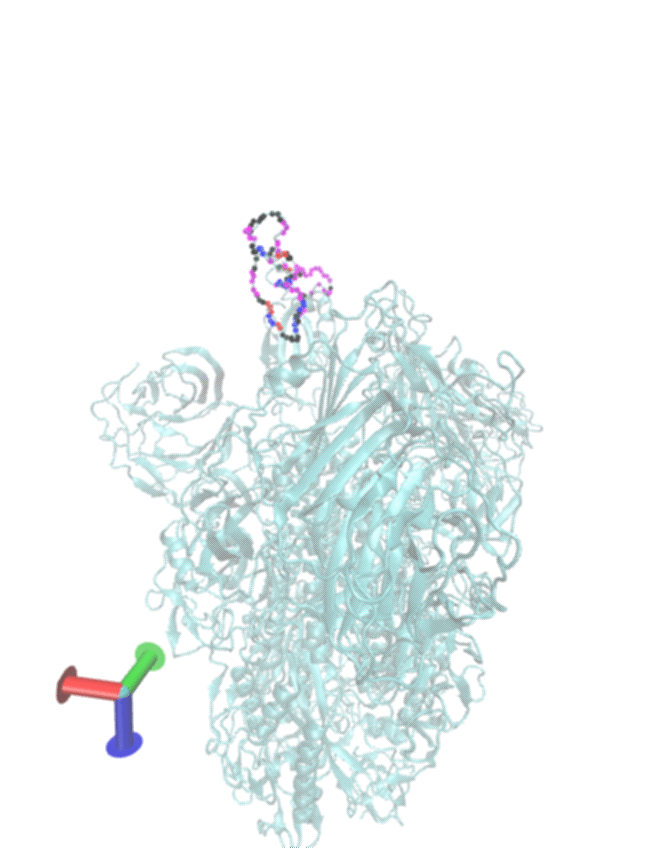

### Supporting Information Movie S.9

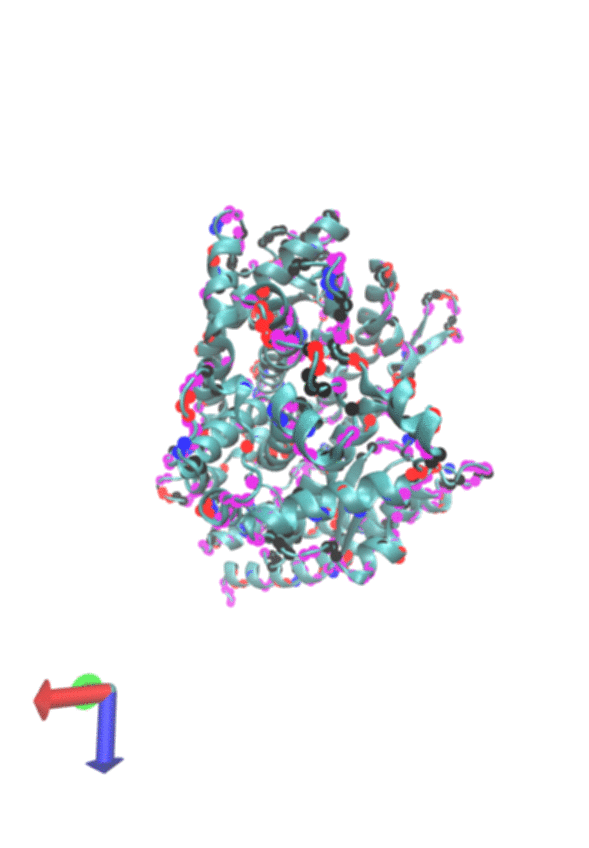

### Supporting Information Movie S.10

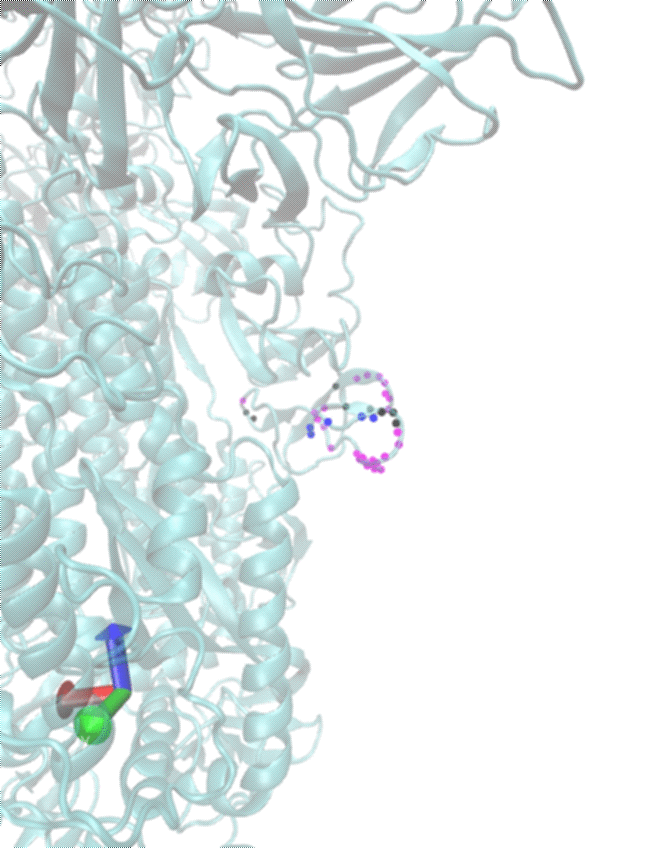

### Supporting Information Movie S.11

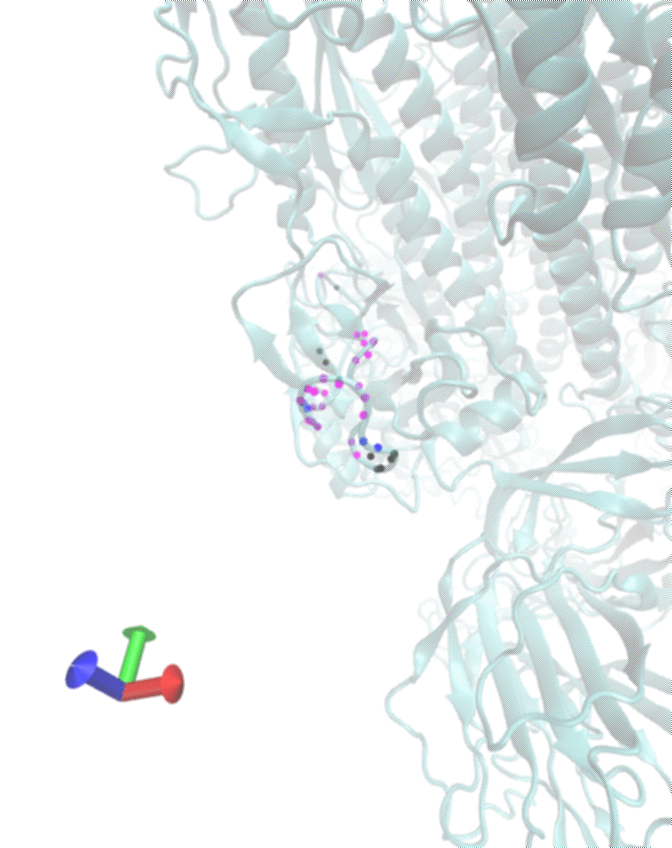
