## Supporting Information for "Protein Surface Printer for Exploring Protein Domains"

### S1. PETase trajectory alignment and multiframe analysis

Table S.1: Surface domains of PETase

| Software/ Force field | Positively charged | Negatively charged | Polar Neutral | Non polar |
|---|---|---|---|---|
| GROMACS CHARMM36m | 6.2% ± 0.5% | 4.9% ± 0.5% | 61.0% ± 0.8% | 27.9% ± 1.0% |
| NAMD CHARMM36 | 6.3% ± 0.5% | 4.9% ± 0.4% | 61.2% ± 1.1% | 27.6% ± 1.0% |
| AMBER AMBER14SB | 6.4% ± 0.6% | 4.9% ± 0.5% | 60.3% ± 1.3% | 28.3% ± 1.1% |
| OpenMM AMOEBA 13 | 6.0% ± 0.5% | 5.2% ± 0.4% | 60.7% ± 1.0% | 28.0% ± 1.0% |

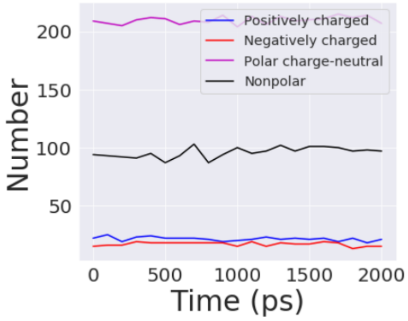


(a)

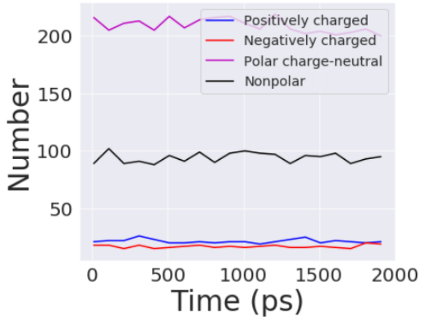


(b)

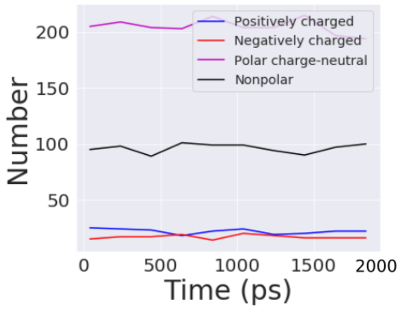


(c)

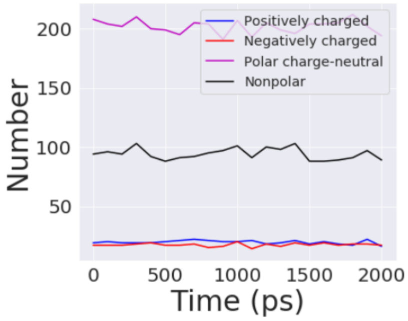


(d)

Fig. S. 1: Percentage of surface domains change over time. Color legends are shown in figures. (a) Gromacs/CHARMM36m (b) NAMD/CHARMM36 (c) AMBER/AMBER14SB (d) OpenMM/AMEOBA 13

#### S2. P450 dimer single frame analysis

Table S.2: Surface domains of P450 dimer

| Positively charged | Negatively charged | Polar Neutral | Non polar |
|---|---|---|---|
| 14.9% | 19.1% | 30.7% | 35.3% |

#### S3. Case study: SARS-CoV-2 and SARS-CoV analysis

MFVFLVLLPLVSSQCVNLTTRTQLPPAYTNSFTRGVYYPDKVFRSSVLHSTQDLFLPFFSNVTWFHAIHVSGTNGTKRFD 1-80

NPVLPFNDGVYFASTEKSNIIRGWIFGTTLDSKTQSLLIVNNATNVVIKVCEFQFCNDPFLGVYYHKNNKSWMESEFRVY 81-160

SSANNCTFEYVSQPFLMDLEGKQGNFKNLREFVFKNIDGYFKIYSKHTPINLVRDLPQGFSALEPLVDLPIGINITRFQT 161-240

LLALHRSYLTPGDSSSGWTAGAAAYYVGYLQPRTFLLKYNENGTITDAVDCALDPLSETKCTLKSFTVEKGIYQTSNFRV 241-320

QPTESIVRFPNITNLCPFGEVFNATRFASVYAWNRKRISNCVADYSVLYNSASFSTFKCYGVSPTKLNDLCFTNVYADSF 321-400

VIRGDEVRQIAPGQTGKIADYNYKLPDDFTGCVIAWNSNNLDSKVGGNYN**YLYRLFRKSNLKPFERDISTEIYQAGSTPC** 401-480

**NGVEGFNCYFPLQSYGFQPTNGVGYQPYR**VVVLSFELLHAPATVCGPKKSTNLVKNKCVNFNFNGLTGTGVLTESNKKFL 481-560

PFQQFGRDIADTTDAVRDPQTLEILDITPCSFGGVSVITPGTNTSNQVAVLYQDVNCTEVPVAIHADQLTPTWRVYSTGS 561-640

NVFQTRAGCLIGAEHVNNSYECDIPI**GAGICASYQTQTNSPRRARSVASQSII**AYTMSLGAENSVAYSNNSIAIPTNFTI 641-720

SVTTEILPVSMTKTSVDCTMYICGDSTECSNLLLQYGSFCTQLNRALTGIAVEQDKNTQEVFAQVKQIYKTPPIKDFGGF 721-800

NFSQILPDPSKPSKRSFIEDLLFNKVTLADAGFIKQYGDCLGDIAARDLICAQKFNGLTVLPPLLTDEMIAQYTSALLAG 801-880

TITSGWTFGAGAALQIPFAMQMAYRFNGIGVTQNVLYENQKLIANQFNSAIGKIQDSLSSTASALGKLQDVVNQNAQALN 881-960

TLVKQLSSNFGAISSVLNDILSRLDKVEAEVQIDRLITGRLQSLQTYVTQQLIRAAEIRASANLAATKMSECVLGQSKRV 961-1040

DFCGKGYHLMSFPQSAPHGVVFLHVTYVPAQEKNFTTAPAICHDGKAHFPREGVFVSNGTHWFVTQRNFYEPQIITTDNT 1041-1120

FVSGNCDVVIGIVNNTVYDPLQPELDSFKEELDKYFKNHTSPDVDLGDISGINASVVNIQKEIDRLNEVAKNLNESLIDL 1121-1200

QELGKYEQYIKWPWYIWLGFIAGLIAIVMVTIMLCCMTSCCSCLKGCCSCGSCCKFDEDDSEPVLKGVKLHYT 1201-1273

RBD

O-linked Glycan residues

Fig. S. 2: The sequence of SARS-CoV-2 spike protein.

| | |
|---|---|
| MFIFLLFLTLTSGSDLDRCTTFDDVQAPNYTQHTSSMRGVYYPDEIFRSDTLYLTQDLFLPFYSNVTGFHTINHTFGNPV | 1-80 |
| IPFKDGIYFAATEKSNVVRGWVFGSTMNNKSQSVIIINNSTNVVIRACNFELCDNPFFAVSKPMGTQTHTMIFDNAFNCT | 81-160 |
| FEYISDAFSLDVSEKSGNFKHLREFVFKNKDGFLYVYKGYQPIDVVRDLPSGFNTLKPIFKLPLGINITNFRAILTAFSP | 161-240 |
| AQDIWGTSAAAYFVGYLKPTTFMLKYDENGTITDAVDCSQNPLAELKCSVKSFEIDKGIYQTSNRVVPSGDVVRFPNITN | 241-320 |
| LCPFGEVFNATKFPSVYAWERKKISNCVADYSVLYNSTFFSTFKCYGVSATKLNDLCFSNVYADSFVVKGDDVRQIAPGQ | 321-400 |
| TGVIADYNYKLPDDFMGCVLAWNTRNIDATSTGNYN**YKYRYLRHGKLRPFERDISNVPFSPDGKPCTPPALNCYWPLNDY** | 401-480 |
| **GFYTTTGIGYQPYR**VVVLSFELLNAPATVCGPKLSTDLIKNQCVNFNFNGLTGTGVLTPSSKRFQPFQQFGRDVSDFTDS | 481-560 |
| VRDPKTSEILDISPCSFGGVSVITPGTNASSEVAVLYQDVNCTDVSTAIHADQLTPAWRIYSTGNNVFQTQAGCLIGAEH | 561-640 |
| VDTSYECDIPI**GAGICASYHTVSLLRSTSQKSIV**AYTMSLGADSSIAYSNNTIAIPTNFSISITTEVMPVSMAKTSVDCN | 641-720 |
| MYICGDSTECANLLLQYGSFCTQLNRALSGIAAEQDRNTREVFAQVKQMYKTPTLKYFGGFNFSQILPDPLKPTKRSFIE | 721-800 |
| DLLFNKVTLADAGFMKQYGECLGDINARDLICAQKFNGLTVLPPLLTDDMIAAYTAALVSGTATAGWTFGAGAALQIPFA | 801-880 |
| MQMAYRFNGIGVTQNVLYENQKQIANQFNKAISQIQESLTTTSTALGKLQDVVNQNAQALNTLVKQLSSNFGAISSVLND | 881-960 |
| ILSRLDKVEAEVQIDRLITGRLQSLQTYVTQQLIRAAEIRASANLAATKMSECVLGQSKRVDFCGKGYHLMSFPQAAPHG | 961-1040 |
| VVFLHVTYVPSQERNFTTAPAICHEGKAYFPREGVFVFNGTSWFITQRNFFSPQIITTDNTFVSGNCDVVIGIINNTVYD | 1041-1120 |
| PLQPELDSFKEELDKYFKNHTSPDVDLGDISGINASVVNIQKEIDRLNEVAKNLNESLIDLQELGKYEQYIKWPWYVWLG | 1121-1200 |
| FIAGLIAIVMVTILLCCMTSCCSCLKGACSCGSCCKFDEDDSEPVLKGVKLHYT | 1201-1254 |

RBD

O-linked Glycan residues

Fig. S. 3: The sequence of SARS-CoV spike protein.

Table S.3: Surface domains of the spike protein.

| Virus | Positively charged | Negatively charged | Polar Neutral | Non polar |
|---|---|---|---|---|
| SARS-CoV-2 | 9.0% | 10.4% | 47.2% | 33.4% |
| SARS-CoV | 9.2% | 11.0% | 45.5% | 34.3% |

Table S.4: Surface domains of RBD.

| Virus | Positively charged | Negatively charged | Polar Neutral | Non polar |
|---|---|---|---|---|
| SARS-CoV-2 | 10.3% | 7.4% | 50.0% | 32.3% |
| SARS-CoV | 17.2% | 8.6% | 39.0% | 35.2% |

Table S.5: Surface domains of ACE2.

| | Positively charged | Negatively charged | Polar Neutral | Non polar |
|---|---|---|---|---|
| Outer Surface | 9.8% | 19.4% | 37.2% | 33.6% |
| Inner Surface | 13.1% | 16.8% | 45.3% | 24.8% |

#### S4. 3D printout of each analysis

Movie S.1: 3D printout of PETase surface from Gromacs

Movie S.2: 3D printout of PETase surface from NAMD

Movie S.3: 3D printout of PETase surface from AMBER

Movie S.4: 3D printout of PETase surface from OpenMM

Movie S.5: 3D printout of SARS-CoV-2 spike protein surface

Movie S.6: 3D printout of SARS-CoV spike protein surface

Movie S.7: 3D printout of SARS-CoV-2 RBD

Movie S.8: 3D printout of SARS-CoV RBD

Movie S.9: 3D printout of ACE2 surface

Movie S.10: 3D printout of SARS-CoV-2 O-linked residues

Movie S.11: 3D printout of SARS-CoV O-linked residues
